# A PMMA bone cement composite that functions as a drug-refillable depot for localized, multi-window chemotherapeutic treatment of bone cancer

**DOI:** 10.1101/2021.06.13.448250

**Authors:** Erika L. Cyphert, Nithya Kanagasegar, Ningjing Zhang, Greg D. Learn, Horst A. von Recum

## Abstract

Standard chemotherapy for primary and secondary bone tumors typically involves systemic administration of chemotherapeutic drugs, such as doxorubicin (DOX). However, non-targeted delivery increases dose requirements, and results in off-target toxicity and suboptimal chemotherapeutic efficacy. When chemotherapy is ineffective, substantial resection of tissue and/or total amputation become necessary – a debilitating outcome for any patient. In this work, we developed a proof-of-concept, non-biodegradable, mechanically robust, and refillable composite system for chemotherapeutic (i.e. DOX) delivery comprised of poly(methyl methacrylate) (PMMA) bone cement and insoluble polymeric γ-cyclodextrin (γ-CD) microparticles. The porosity and compressive strength of DOX-filled PMMA composites were characterized. DOX filling capacity, elution kinetics, cytotoxicity against primary osteosarcoma and lung cancer cells, and refilling capacity of composites were evaluated. PMMA composites containing up to 15wt% γ-CD microparticles provided consistent, therapeutically-relevant release of DOX with ~100% of the initial DOX released after 100 days. Over the same period, only ~6% of DOX was liberated from PMMA with free DOX. Following prolonged curing, PMMA composites with up to 15wt% γ-CD surpassed compressive strength requirements outlined by international standards for acrylic bone cements. Compared to DOX-filled PMMA, DOX-filled PMMA/γ-CD composites provided long-term release with decreased burst effect, correlating to long-term cytotoxicity against cancer cells. Refillable properties demonstrated by the PMMA composite system may find utility for treating local recurrences, limiting chemoresistance, and altering drug combinations to provide customized treatment regimens. Overall, findings suggest that PMMA composites have the potential to serve as a platform for the delivery of combinatorial chemotherapeutics to treat bone tumors.

## Introduction

Bone tumors account for approximately 1% new cancer diagnoses annually in the United States and are most prevalent among children and adolescents, representing the 3^rd^ most common childhood malignancy [1]–[3]. Despite their relatively low overall rate of occurrence, primary bone-derived tumors can be particular harrowing to treat; their necessary resection results in considerable morbidity (e.g. bone defects) and amputations become necessary in up to 10% of patients [1]. Certain types of bone tumors, such as malignant osteosarcomas and Ewing sarcomas can result in metastatic lung lesions, further complicating treatment and decreasing remission and 5-year survival rates (e.g. 20-30% survival rate versus 70-80% survival rate with localized lesion) [1], [4]–[7]. Additionally, several other types of cancer including breast [8]–[10], lung adenocarcinoma [1], [11], [12], and prostate [13] have been shown to recur as bone metastases. It has been reported that lung cancer contributes to between 30-70% of bone metastases [11]. Survival rates in cancer patients with bone metastases dramatically decline as metastases typically present in the late stages of cancer (i.e ~60% decrease in 5-year survival rate) [4]–[7]. Bone metastases are also responsible for considerable pain and are challenging to locate and treat in a targeted manner [10]–[12].

Treatment of localized bone tumors generally involves a multi-pronged approach consisting of systemic neoadjuvant chemotherapy (doxorubicin – DOX, cisplatin, methotrexate) followed by surgical resection and adjuvant chemotherapy [1], [14], [15]. To effectively reduce the size of the tumor lesion prior to excision, high systemic doses of chemotherapeutics are often required [1], [15]. To prevent tumor recurrence, proper excision of tumor margins is essential; however, due to challenges of differentiating between healthy and cancerous tissue, and the serious health risks that recurrence entails, it has historically been necessary for surgeons to excise large sections of bone tissue to ensure total resection [1], [16], [17]. Consequently, surgery often results in large bone defects which can be challenging to reconstruct/regenerate and can inflict substantial mobility restrictions and aesthetic concerns to patients depending upon their location [16], [17]. Bone metastases can be more elusive to treat as they often require whole-body scintigraphy imaging with radioactive tracers to detect, and depending upon the extent of their spread, they may be impossible to surgically excise [18]. Imaging can be used to monitor bone metastases, but treatments in most cases are only palliative to help to reduce pain and include a combination of chemotherapy, radiation therapy, and bone-targeting therapies (bisphosphonates, denosumab – RANKL inhibitor, cathepsin K inhibitors) [10], [19]–[21].

Systemic administration of chemotherapeutics is suboptimal due to the widespread off-target toxicities (i.e. cardio-, hepato-, nephron-toxicity) that can result from prolonged exposure to DOX, cisplatin, and methotrexate, for example [22]–[24]. Furthermore, chemoresistance can develop if doses are administered at sub-therapeutic levels locally, which subsequently complicates treatment regimens and worsens prognosis [25]–[27]. Specifically, usage of DOX, cisplatin, and methotrexate have been shown to contribute to the overexpression of membrane bound drug transporter protein P-glycoprotein [25], upregulation of heat shock protein (HSP90AA1) [26], and high mobility group box 1 (HMGB1)-mediated autophagy [27], driving chemoresistance in bone tumors. As a result, a system capable of delivering chemotherapeutics in a controlled, consistent, and localized manner at therapeutic levels would be ideal to treat primary and secondary bone tumors more effectively, and to reduce chemoresistance, improving patient outcomes.

To reduce the off-target toxicities associated with prolonged systemic administration of chemotherapeutics to treat primary bone tumors [28]–[38] and bone metastases [39]–[46], a variety of localized, site-specific delivery systems have been developed. Specifically, in the treatment of primary tumors, several groups have developed injectable stimuli-responsive biodegradable polymeric carriers for chemotherapeutic agents such as DOX, cisplatin, methotrexate, and gemcitabine comprised of poly(lactic-co-glycolic acid) (PLGA), poly(ethylene glycol) (PEG), poly(*N*-isopropylacrylamide-co-acrylamide), and gelatin methacryloyl [28], [34]–[36], [38]. Other systems have utilized bioactive materials such as hydroxyapatite nanoparticles and mesoporous bioactive glass to promote bone formation while eradicating tumor cells [31], [37]. For localized treatment of bone metastases, various chemotherapeutic-filled nanoparticles and micelles have been developed to target overexpressed CD44 tumor antigen or various osteoclastic pathways (i.e. bone-targeted glutamic oligopeptides, Gli antagonist – GANT58) to locally deliver the payload [39], [40], [42], [44]–[46]. While these systems have been shown to help eradicate tumor cells and reduce the size of the primary tumor both *in vitro* and *in vivo*, they generally have a short therapeutic window of activity. Many systems degrade within 30 days and only exhibit release of the chemotherapeutic agent for a couple of days.

As an alternative to injectable and biodegradable localized chemotherapeutic delivery systems, poly(methyl methacrylate) (PMMA) bone cement has been utilized as a depot to treat bone tumors [47]–[51]. Due to its prior longstanding use in orthopaedic surgery for fixation of prostheses during arthroplasties, PMMA bone cement was one of the first vehicles used for localized delivery of antibiotics in the treatment of infections involving bone (namely periprosthetic joint infection and osteomyelitis) [52]. PMMA thus represents an obvious candidate as a vehicle material for chemotherapeutic delivery. However, drug release from PMMA is severely limited for several reasons. First, many encapsulated compounds have reduced activity following heat exposure from the exothermic cement polymerization process. Second, for the drug that is initially mixed into the cement, most remains permanently entrapped in the cement, with >90% remaining stuck after 90 days [47], with the remainder being released essentially all at once shortly after implantation (i.e. burst release). This results in sub-therapeutic levels of drug reaching the tumor cells, promoting chemoresistance.

Currently, there are no bone tumor chemotherapeutic delivery systems capable of providing long-term controlled and localized release that can be refilled on-demand and in a patient-customized manner. A potential solution to this unmet need may be inspired by a solution we have previously developed to enable controlled delivery of antibiotics using PMMA bone cement using insoluble cyclodextrin (CD) microparticles. CD is a cyclic oligosaccharide comprised of 6-8 glucose units (α-, β-, γ-), forming a toroid structure with a slightly hydrophobic pocket. Small molecule drugs can be entrapped and released from CD pockets based upon their binding strength or affinity [53], [54]. We have previously incorporated polymerized insoluble CD microparticles into PMMA bone cement, and the composite system has demonstrated its capacity to be repeatedly refilled with antibiotics after implantation and to release the drugs in a controlled and consistent manner [55]–[58]. Incorporation of 10wt% CD microparticles in the PMMA did not negatively impact its ultimate compressive strength relative to clinically-used controls [55], [56], [58]. Here, we incorporated γ-CD microparticles into PMMA for the delivery of DOX due to the strong affinity between DOX and γ-CD (−5.98 kcal) [59]. Moreover, the ability to refill the PMMA composite with chemotherapeutics after implantation is advantageous because it theoretically would enable surgeons to customize the patient’s treatment and locally administer an additional bolus of the initial drug or a different drug if the tumor is to recur. In existing systems, addition of drugs following implantation is not possible.

In this study, we have developed a proof-of-concept PMMA composite system for the controlled delivery of DOX comprised of different amounts of γ-CD microparticles (i.e. 10wt% and 15wt%). The relative porosity and ultimate compressive strength of the PMMA composites were evaluated to determine if the composites could be utilized in load-bearing applications. Additionally, we compared the release kinetics and cytotoxicity of collected DOX aliquots from PMMA with and without γ-CD microparticles against both osteosarcoma (MG-63) and lung carcinoma (Lewis Lung Carcinoma – LLC) cell lines using 3-(4,5-dimethylthiazol-2-yl)-5-(3-carboxymethyoxyphenyl)-2-(4-sulfophenyl)-2H-tetrazolium (MTS) studies to simulate the treatment of primary osteosarcoma lesions and lung-derived bone metastases [1], [11], [12]. Finally, agarose-based tissue-mimicking refilling models were utilized to explore the ability of PMMA composites to be refilled with DOX following implantation. **Figure 1** depicts an overview of the analyses carried out in this work.

**Figure 1.**
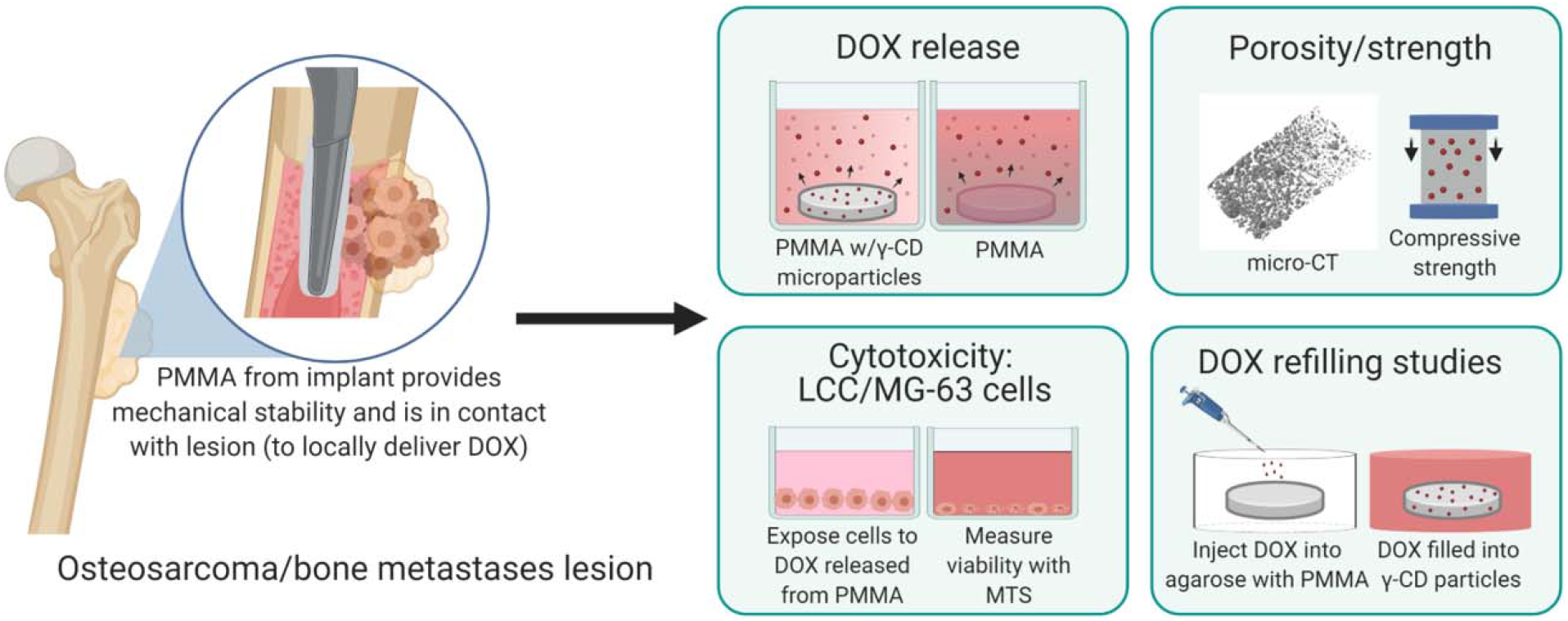
Schematic overview of work. Primary osteosarcoma or bone metastases lesions have the potential to be treated via localized chemotherapeutic delivery from poly(methyl methacrylate) (PMMA) bone cement. PMMA bone cement composites were prepared through the incorporation of different amounts of cyclodextrin (γ-CD) microparticles with doxorubicin (DOX). PMMA composites were evaluated for their A) DOX release kinetics, B) cytotoxicity against Lewis Lung Carcinoma (LLC) and osteosarcoma (MG-63) cells, C) porosity and ultimate compressive strength, and D) their ability to be refilled with DOX after implantation. Figure created using BioRender.

## Experimental

### Materials

Lightly epichlorohydrin cross-linked γ-CD pre-polymer was purchased from CycloLabs (Budapest, Hungary). Ethylene glycol diglycidyl ether was purchased from Polysciences Inc. (Warrington, PA). Doxorubicin hydrochloride (DOX) was purchased from LC Laboratories (Woburn, MA). Simplex^®^ HV (high viscosity) radiopaque bone cement (Stryker Orthopaedics, Mahwah, NJ) was obtained from eSutures (Mokena, IL). Lewis Lung Carcinoma (LLC) cells and MG-63 human osteosarcoma cells were purchased from ATCC^®^ (Manassas, VA) (ATCC^®^ CRL-1642™, ATCC^®^ CRL-1427™). Dulbecco’s Modified Eagle’s Medium – high glucose (DMEM), fetal bovine serum (FBS), and penicillin/streptomycin were obtained from Gibco (Carlsbad, CA). All other reagents were purchased in the highest grade possible from Fisher Scientific.

### Cyclodextrin (CD) microparticle synthesis

Insoluble gamma-cyclodextrin (γ-CD) microparticles were prepared according to a previously published protocol [55]–[58], [60]. Briefly, 1 gram of epichlorohydrin cross-linked γ-CD was dissolved in 4 mL of 0.2 M potassium hydroxide and combined with 1.6 mL ethylene glycol diglycidyl ether for cross-linking. The mixture was poured into 50 mL light mineral oil with 750 μL surfactant (24% Tween 85, 76% Span 85), and stirred and heated at 60°C for 4 hours. The resulting γ-CD microparticles were washed and centrifuged twice in 35 mL of each solvent: light mineral oil, hexanes, acetone, and MilliQ water. γ-CD microparticles were frozen, lyophilized and stored in a desiccator until further use.

### Loading CD microparticles with doxorubicin (DOX)

20 mg samples of dried γ-CD microparticles were placed in 1 mL of DOX dissolved in dimethyl sulfoxide (DMSO) (5 mg/mL), covered in foil, and placed on a rotisserie shaker for 72 hours [55]–[58]. DOX-filled γ-CD microparticles were washed 5x with MilliQ water, frozen, lyophilized, and stored in a desiccator until further use.

### PMMA fabrication (beads and cylinders)

PMMA fabrication was carried out in small batches according to the manufacturer’s instructions [55]–[58]. Specifically, for each batch, 2 grams of Simplex^®^ HV surgical grade bone cement powder was combined with 1 mL of methyl methacrylate monomer and mixed with the desired additives (i.e. DOX, γ-CD microparticles, or DOX-filled γ-CD microparticles). For DOX control samples (without γ-CD microparticles), batches containing free 5 mg (1.25% wt/wt DOX/PMMA) were prepared. For samples containing CD microparticles, either 5, 10, or 15wt% empty (non-drug filled) or DOX-filled γ-CD microparticles were added to the PMMA powder prior to polymerization and mixed until homogeneous. Once a soft dough formed, it was either rolled into a thin layer (~ 2 mm thickness) and punched into small beads (6 mm diameter) or finger pressed into 12 mm height × 6 mm diameter cylindrical molds and cured at room temperature.

### Micro-computed tomography (micro-CT)

To determine the porosities of PMMA samples with DOX and DOX-filled γ-CD microparticles, micro-computed tomography (micro-CT) was used. Settings for the micro-CT scans were based upon a published methodology [55], [56], [58]. Specifically, PMMA cylinders were scanned inside of polypropylene tubes using a Siemens Inveon PET-CT scanner (Siemens Medical Solutions, Malvern, PA) controlled by Siemens Inveon Acquisition Workplace software on a PC. To ensure consistency, past scanning and reconstruction settings were utilized for each sample (i.e. medium-high magnification, 80 kV, 500 μA, binning factor = 1, 448 ms exposure, 200 ms settle, noncontinuous rotation, 512 steps, 360° rotation, 14.21 μm pixel size) [55], [56], [58]. For segmentation of the pores, data was exported in DICOM format to 3D Slicer software (BWH and 3D Slicer contributors, version 4.8) [61] with an upper threshold of −200 HU. Volumetric measurements (i.e. number of pores and pore volume fraction) were calculated using Netfabb Standard 2020 (Autodesk, Inc.) using models exported from 3D Slicer. Scans were collected on three experimental PMMA groups in triplicate: free 1.25% DOX (no γ-CD), 10wt% DOX-filled γ-CD microparticles, and 15wt% DOX-filled γ-CD microparticles.

### Mechanical compression studies

To evaluate the impact of additives (i.e. free DOX and DOX-filled γ-CD microparticles) on the mechanical strength of PMMA composites, compressive strength studies were carried out. Parameters used for compressive strength testing were determined based on previously published methodologies [55], [56], [58]. Specifically, the ends of 12 mm × 6 mm PMMA composite cylinders were sanded square using a drill press and wet 240 grit silicon carbide sandpaper. The mass, diameter, and length of each machined cylinder was measured using a digital scale and calipers. To be included in the study, samples were visually inspected to ensure that they did not contain defects > 0.5 mm in diameter. ASTM 451-16 standards were used to guide the set-up of the compressive testing where cylinders were loaded under unconfined compression at a rate of 20 mm/min (adjusted to 19 mm/min for slightly shorter cylinders) and sample rate of 200 Hz with a mechanical testing frame (Material Testing System MTS 810, MTS Systems Corporation) with a 200 lbf (8896 N) load cell [62]. To evaluate the impact of the duration of time between curing of the PMMA cylinders and mechanical testing on their resultant compressive strength, cylinders were tested at both 48 hours (plain controls, free 1.25% DOX, 15wt% DOX-filled γ-CD microparticles) and 4-5 months following fabrication (free 1.25% DOX, 10wt% and 15wt% DOX-filled γ-CD microparticles). Ultimate compressive strength, modulus, and normalized work to failure were calculated from load-displacement curves and sample dimensions. 2% offset was used to determine the ultimate compressive strength (MPa) of each sample. Each condition was evaluated with a minimum of 3-6 replicates.

### Agarose refilling model

To evaluate the ability of PMMA containing γ-CD microparticles to be refilled with DOX following implantation in tissue, an agarose-based tissue-mimicking refilling model was used. Agarose gels were prepared as previously described [55]–[58], [63], where 5 mL of hot agarose dissolved in phosphate buffered saline (PBS) (0.075% wt/vol) was placed into each well of a 6-well plate. A small PMMA bead (plain: without drug or γ-CD; 5wt%, 10wt%, or 15wt% empty γ-CD microparticles: without drug) was placed on top of the solidified agarose and an additional 5 mL of hot agarose was placed on top to embed the PMMA bead. Once the gel set, a 4 mm diameter well was punched into the center of the gel and 100 μL of 5 mg/mL DOX dissolved in DMSO was injected into the well. The plate was covered in foil and placed into a shaking incubator (37°C) for 48 hours to allow for the drug to diffuse into the embedded PMMA. Following incubation, PMMA beads were removed from the agarose and dried for future analysis Each condition was carried out in triplicate.

### Quantification of filling and refilling capacity

To quantify the mass of DOX initially filled (i.e. prefilled) and refilled into PMMA beads in the agarose-based model, PMMA samples were dissolved in DMSO to leach-out the DOX [55], [56]. Specifically, individual PMMA beads were placed in 3 mL of DMSO, covered in foil, and placed in a shaking incubator at 37°C for 48 hours. Once PMMA beads had dissolved, 200 μL aliquots of the solution were placed into each well of a 96-well plate and the mass of DOX in the dissolved solution was calculated using a calibration curve and by measuring the fluorescence signal (excitation = 498 nm, emission = 590 nm) using a Biotek™ 96-well plate reader (H1: Winooski, VT).

### Fluorescence imaging and quantification

To quantify and image the relative fluorescence intensity of the DOX filled into the surface of the PMMA beads, *in vivo* fluorescence imaging was used. PMMA beads containing DOX (free 1.25% DOX, pre- and re-filled γ-CD microparticles with DOX) were placed inside a Maestro™ *In-Vivo* Fluorescence Imaging System (Cambridge Research & Instrumentation, Inc. Cri, Woburn, MA) with the following settings: filter = green, stage height/lights = 2B, exposure = 47.33 ms. For fluorescence intensity quantification, PMMA beads were aligned in triplicate, autofluorescence signal was removed, and uniform regions of interest were drawn (5 mm diameter). The average total fluorescence signal (RFU) generated from DOX in PMMA samples was normalized by the signal from the PMMA samples of the same composition without DOX.

### DOX release

To determine the release kinetics of DOX from different PMMA compositions, release aliquots were collected at discrete time points in PBS supplemented with a hydrophobic sink (Tween 80). Specifically, individual PMMA beads (prefilled and refilled with DOX) were weighed and placed into a 24-well plate in 1 mL of PBS with 5% Tween 80 to provide a hydrophobic sink for the DOX complexed with γ-CD to be released. Plates were covered in foil and placed in a shaking incubator (37°C). After set time points (i.e. 2, 24, 48, 96 hours, etc.) the entire release solution was removed and replaced with fresh buffer to simulate infinite sink conditions. The mass of DOX released at each time point was determined using calibration curves and measuring the fluorescence signal (excitation = 498 nm, emission = 590 nm) using a Biotek™ plate reader. Each condition was completed in triplicate.

### Cell culture

Human osteosarcoma (MG-63) cells and Lewis Lung Carcinoma (LLC) cells were cultured in Dulbecco’s Modified Eagle’s Medium with high glucose and supplemented with 10% fetal bovine serum (FBS) and 1% penicillin/streptomycin at 37°C with 5% CO_2_. Cell utilized in cytotoxicity studies were passaged 3-4 times upon thawing prior to use in studies. MG-63 and LLC cells were passaged over 2 days and 3-4 days, respectively, until cells were confluent.

### MTS cytotoxicity studies

To ensure that the amount of DOX released from prefilled PMMA composites was therapeutically sufficient to eradicate tumor cells over time, cytotoxicity studies were performed. MTS [3-(4,5-dimethylthiazol-2-yl)-5-(3-carboxymethoxyphenyl)-2-(4-sulfophenyl)-2H-tetrazolium] cytotoxicity studies were completed using DOX release aliquots in sterile DMEM (high glucose, 1% penicillin/streptomycin, no FBS) after 1, 7, and 14 days.

Specifically, the cumulative amount of DOX released after 1 day, between 1-7 days, and between 7-14 days was placed on the cells. A 100 μL volume of cell suspension was seeded at either a concentration of 170,000/mL (for MG-63 cells) or 200,000/mL (for LLC cells) in each well in a 96-well plate and were incubated at 37°C (5% CO_2_) for 48 hours. MG-63 and LLC cells were grown and incubated in separate plates. The following controls were included in each plate: blank – no cells, no DOX; growth – cells, no DOX. Following incubation, media was replaced with 100 μL of DOX in sterile DMEM, corresponding to the concentration at each time point (previously determined in drug release studies) in accordance with the values reported in **Table 1** and incubated for 24 hours (37°C, 5% CO_2_) (MG-63 and LLC) or 48 hours (MG-63). Release media was then replaced with 100 μL fresh DMEM (high glucose, 1% penicillin/streptomycin, no FBS) and 20 μL MTS solution. The MTS reaction was allowed to occur over 2 hours at 37°C and 5% CO_2_. After 2 hours, 25 μL of 10% w/v sodium dodecyl sulfate (SDS) was added to block the reaction and solutions were transferred to new 96-well plates and the absorbance was read at 490 nm using Biotek™ plate reader. 6-9 replicates of each condition were plated. The percentage viability following DOX treatment for each well was calculated by subtracting off the background signal from blank controls and then dividing by the growth control.

**Table 1.** Concentration of DOX added to MG-63 and LLC cells after 48 hours of incubation after seeding corresponding to the cumulative amount released from PMMA composite samples after 1 day, between 1 and 7 days, and between 7 and 14 days.

| Time (days) | Free 1.25% DOX<br>(mg/mL) | 10wt% γ-CD (prefilled)<br>(mg/mL) | 15wt% γ-CD (prefilled)<br>(mg/mL) |
| --- | --- | --- | --- |
| 1 | 0.02 | 0.0008 | 0.003 |
| 1-7 | 0.003 | 0.0004 | 0.002 |
| 7-14 | 0.0009 | 0.0003 | 0.0005 |

### Statistical analysis

Data throughout is reported as the average of samples (n= 3) with the exception of compressive studies (Plain PMMA 48 hrs: n = 6, free 1.25% DOX PMMA 48 hrs: n = 3, 15wt% DOX-filled γ-CD PMMA 48 hrs: n = 3, free 1.25% DOX PMMA 4-5 months: n = 4, and 10wt% and 15wt% DOX-filled γ-CD PMMA 4-5 months: n = 4) and cytotoxicity studies (n = 6-9) with the standard deviation as the error bars. Statistical analyses were completed in Microsoft Excel 2016. Single-factor ANOVA with Tukey’s post hoc tests (α = 0.05) were used to analyze micro-CT, mechanical, fluorescence, DOX filling, and cytotoxicity data.

## Results and discussion

### Micro-CT porosity analysis

To visualize the relative distribution of the pores in the different compositions of DOX-filled PMMA, cylindrical samples were scanned using micro-CT. Pores were segmented out from the solid fraction using 3D Slicer and exported to Netfabb for further analysis. Representative 3D renderings of the pore fraction of cylinders without γ-CD microparticles (free 1.25% DOX) and 10wt% and 15wt% DOX-filled γ-CD microparticles are displayed in **Figure 2**.

**Figure 2.**
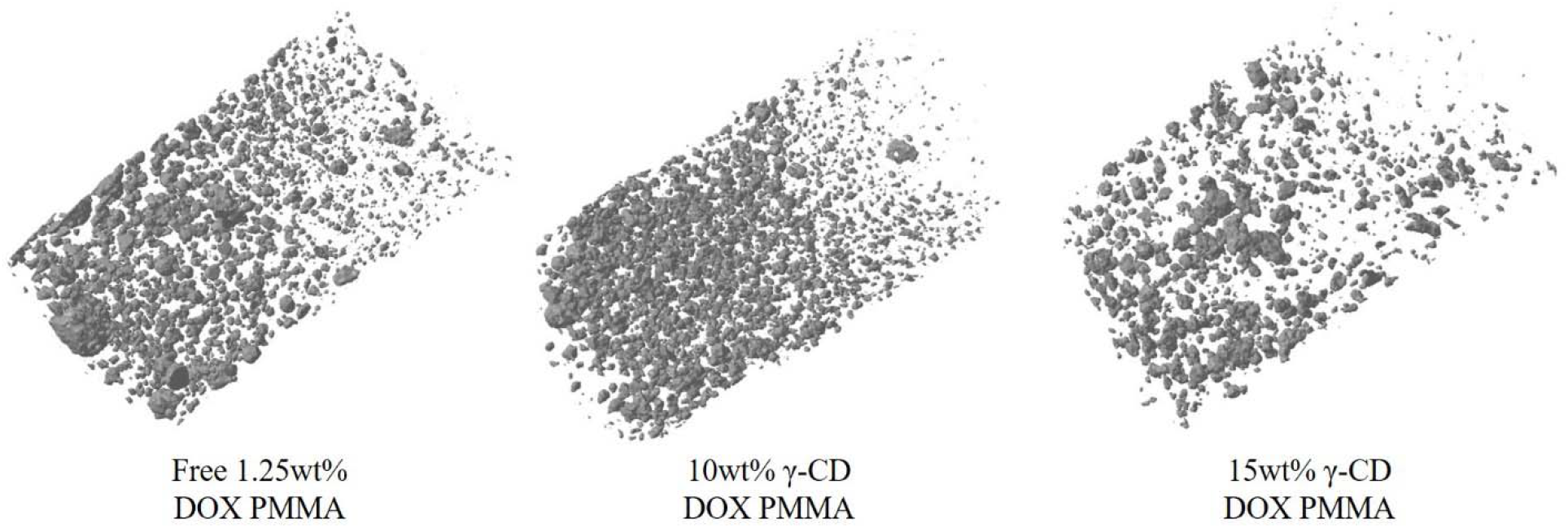
Representative 3D renderings of the segmented pore fraction of free 1.25% DOX (left), 10wt% DOX-filled γ-CD (middle), and 15wt% DOX-filled γ-CD PMMA (right) obtained through micro-CT.

**Figure 3** provides a quantification of several aspects of the porosity including the pore volume fraction (a), average pore volume (b), and average number of discrete pores (c). Interestingly, PMMA containing 15wt% DOX-filled γ-CD microparticles had nearly half the pore volume fraction of samples containing 10wt% γ-CD microparticles and samples without γ-CD microparticles (15wt% = 1.50 ± 0.72%, 10wt% = 2.98 ± 1.17%, free 1.25% DOX = 2.63 ± 0.35%) but not statistically significant. Alternatively, PMMA containing 10wt% DOX-filled γ-CD microparticles had a comparable pore volume fraction to samples containing free 1.25% DOX. In terms of the average volume of individual pores in the samples, sample conditions with 1.25% DOX and 10wt% γ-CD microparticles had similar volumes (free 1.25% DOX = 1.77 ± 0.43 nL, 10wt% γ-CD = 1.62 ± 0.76 nL), whereas samples with 15wt% microparticles had the largest volume (15wt% γ-CD = 2.68 ± 1.59 nL; not statistically significant). PMMA with 10wt% γ-CD microparticles had the largest number of discrete pores (4822 ± 393, p < 0.05) relative to free 1.25% DOX (3851 ± 458) and over 3-times more pores than PMMA with 15wt% γ-CD microparticles (1484 ± 276).

**Figure 3.**
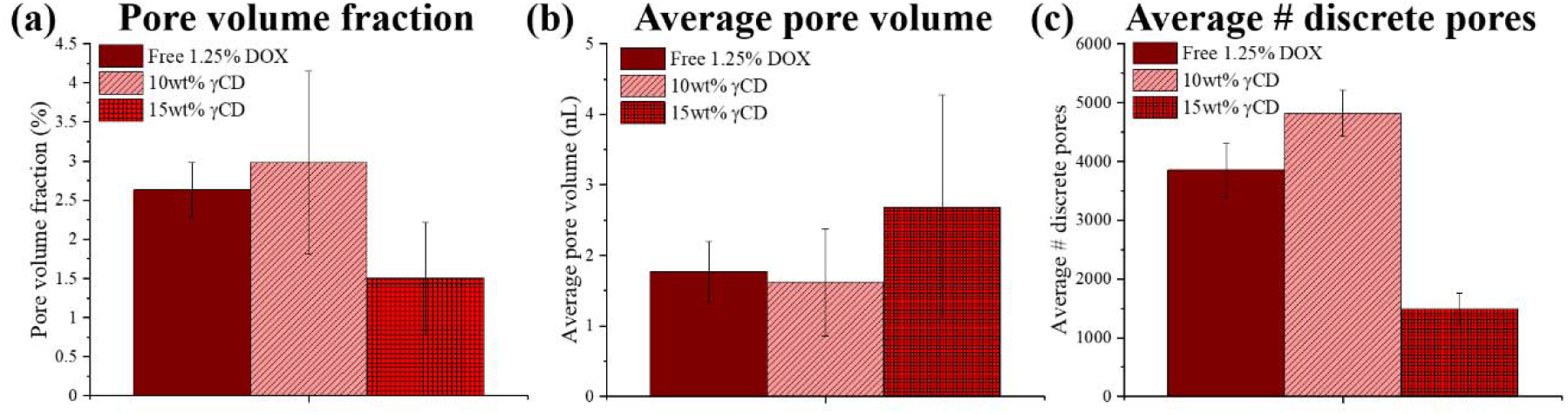
Micro-CT quantification of porosity: (a) pore volume fraction, (b) average pore volume, and (c) average number of discrete pores per sample. PMMA samples were scanned in triplicate with the following compositions: free 1.25% DOX (no γ-CD), 10wt% DOX-filled γ-CD, and 15wt% DOX-filled γ-CD.

Results from the micro-CT analysis demonstrated that pores tended to be aggregated towards the top of the PMMA samples at the PMMA-air interface, as the PMMA undergoes shrinkage during drying [64]. PMMA with 15wt% γ-CD microparticles had fewer but larger pores than those in PMMA with free 1.25% DOX or with 10wt% γ-CD microparticles. Since micro-CT analysis of PMMA composites from our prior studies have indicated that pores are co-registered with γ-CD microparticles due to the radiodensity of the CD polymer (−400 HU) and the upper pore segmentation threshold (−200 HU) [55], [58], it is possible that the γ-CD microparticles in PMMA with 15wt% γ-CD may have clumped, resulting in larger pores. PMMA with 10wt% γ-CD or free 1.25% DOX appeared to have a more even distribution of the pores. Interestingly, PMMA with 15wt% γ-CD had the smallest pore volume fraction. The underlying mechanisms resulting in the decreased porosity in PMMA with 15wt% γ-CD is unknown at this time, but it may be attributed to an interaction between the γ-CD microparticles and the pores. However, since the radiodensity of γ-CD cannot be yet determined following implantation in the PMMA, it is challenging to segment out the γ-CD microparticles from the pores to further probe this phenomenon.

### Mechanical compressive strength analysis

The impact of adding free 1.25% DOX and 10wt% and 15wt% DOX-filled γ-CD microparticles to PMMA on the resultant compressive strength of the PMMA was evaluated. **Figure 4a** depicts representative images of the machined PMMA composite cylinders (12 mm height × 6 mm diameter) that were used in the compressive testing (**Figure 4b**).

**Figure 4.**
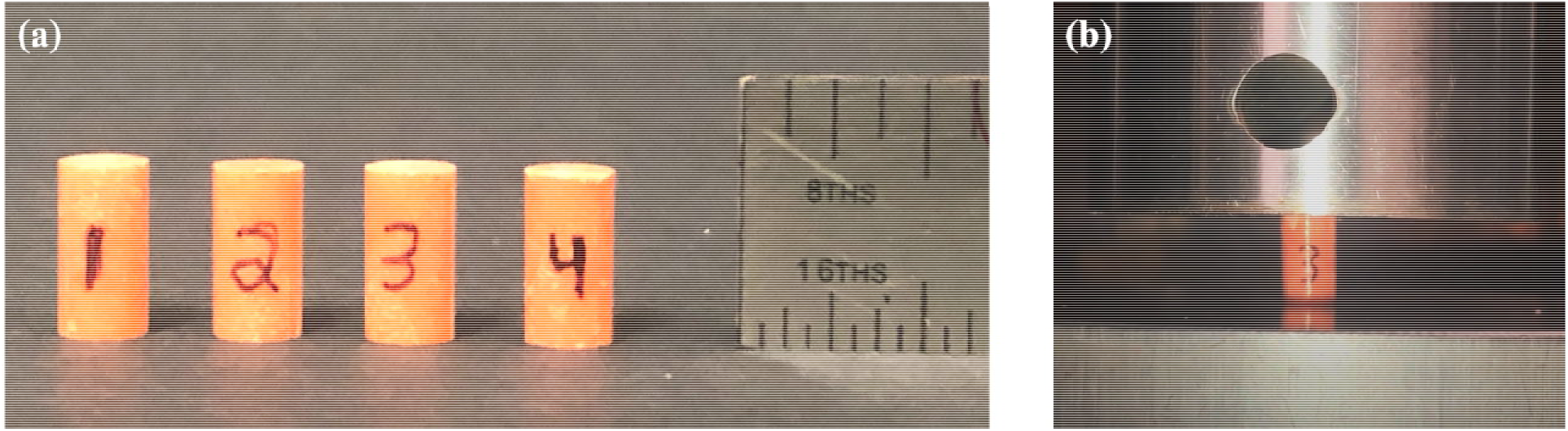
(a) Representative machined PMMA composite cylinders (12 mm height × 6 mm diameter) used in compressive testing. (b) General set-up of compressive testing with machined PMMA composite cylinders.

The effect of the duration of time between the fabrication (curing) of the PMMA composite cylinders and mechanical testing on the compressive strength was evaluated. Specifically, cylinders were fabricated and subsequently mechanically evaluated after either 48 hours or 4-5 months of curing. **Figure 5** depicts representative stress versus strain curves of PMMA composites compressed 48 hours (a) or 4-5 months (b) after curing. 48 hours post-fabrication, the ultimate compressive strength of plain PMMA and PMMA with free 1.25% DOX was significantly greater than PMMA with 15wt% γ-CD microparticles (Plain = 81.4 ± 3.2 MPa, Free 1.25% DOX = 78.5 ± 1.7 MPa, 15wt% γ-CD = 63.4 ± 0.3 MPa). Interestingly, after 4-5 months of curing, there was a statistically significant 24-28% increase in the ultimate compressive strength of PMMA with free 1.25% DOX and 15wt% DOX-filled γ-CD relative to the strength 48 hours post-fabrication (Free 1.25% DOX = 22.4 MPa increase, 15wt% γ-CD = 15.4 MPa increase).

**Figure 5.**
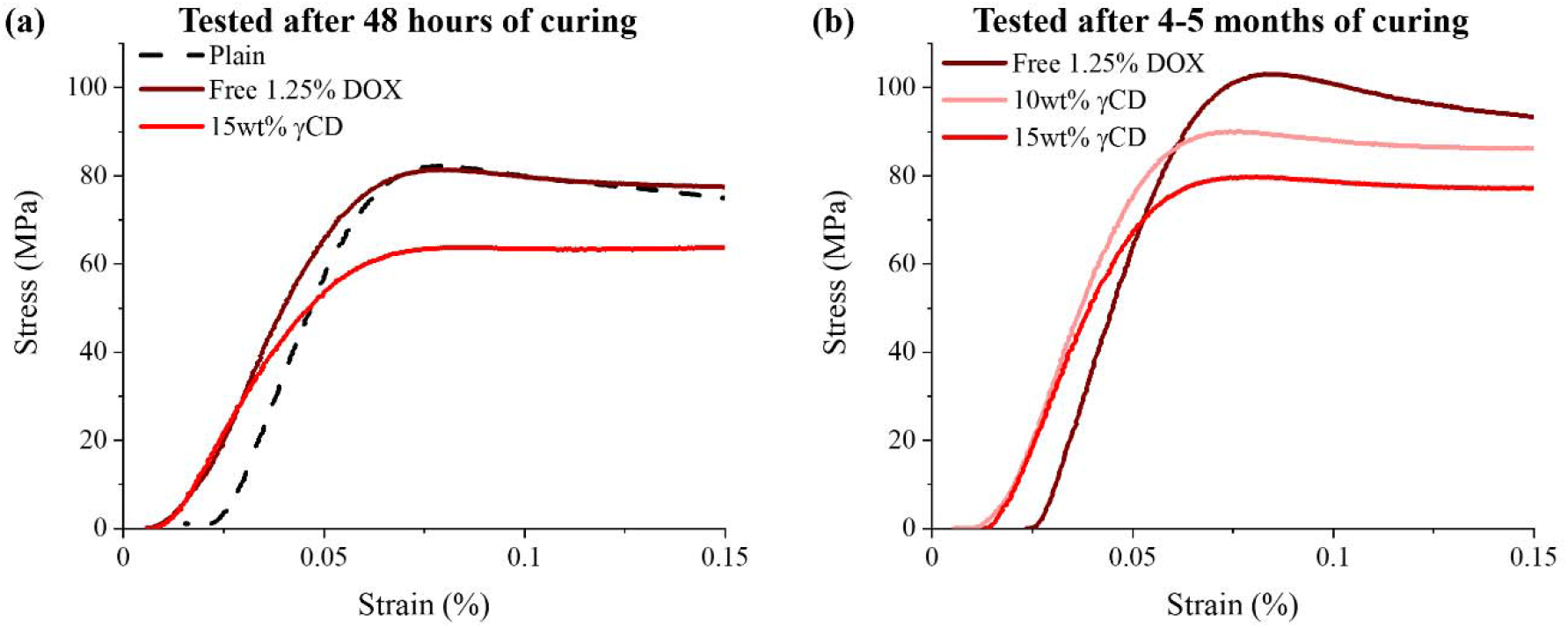
Representative stress versus strain curves of PMMA composite cylinders evaluated at either 48 hours (a) or 4-5 months (b) post-fabrication. Sample conditions included: plain controls (no DOX, no γ-CD), free 1.25% DOX, and 10wt% and 15wt% DOX-filled γ-CD microparticles in PMMA.

**Figure 6** provides a quantification of the ultimate compressive strength (a), modulus (b), and normalized work to failure (c) of PMMA composite cylinders both 48 hours and 4-5 months post-fabrication. 48 hours post-fabrication, plain PMMA had a significantly greater modulus relative to PMMA with 15wt% DOX-filled γ-CD microparticles, but comparable modulus to PMMA with free 1.25% DOX (Plain = 2307.2 ± 159.1 MPa, Free 1.25% DOX = 2044.8 ± 302.1 MPa, 15wt% γ-CD = 1739.5 ± 177.6 MPa). After 4-5 months of curing, there was a statistically significant 33% increase in the modulus of PMMA with free 1.25% DOX and 15wt% DOX-filled γ-CD relative to the strength 48 hours post-fabrication (Free 1.25% DOX = 680.6 MPa increase, 15wt% γ-CD = 588.7 MPa increase). In terms of normalized work to failure, there was no significant difference between the groups 48 hours post-fabrication (Plain = 2.20 ± 0.27 J/cm^3^, Free 1.25% DOX = 2.48 ± 0.17 J/cm^3^, 15wt% γ-CD = 2.47 ± 0.19 J/cm^3^). After 4-5 months of curing, there were no significant increased in the normalized work to failure relative to the values 48 hours post-fabrication (Free 1.25% DOX = 0.91 J/cm^3^ increase, 15wt% γ-CD = 0.37 J/cm^3^ increase).

**Figure 6.**
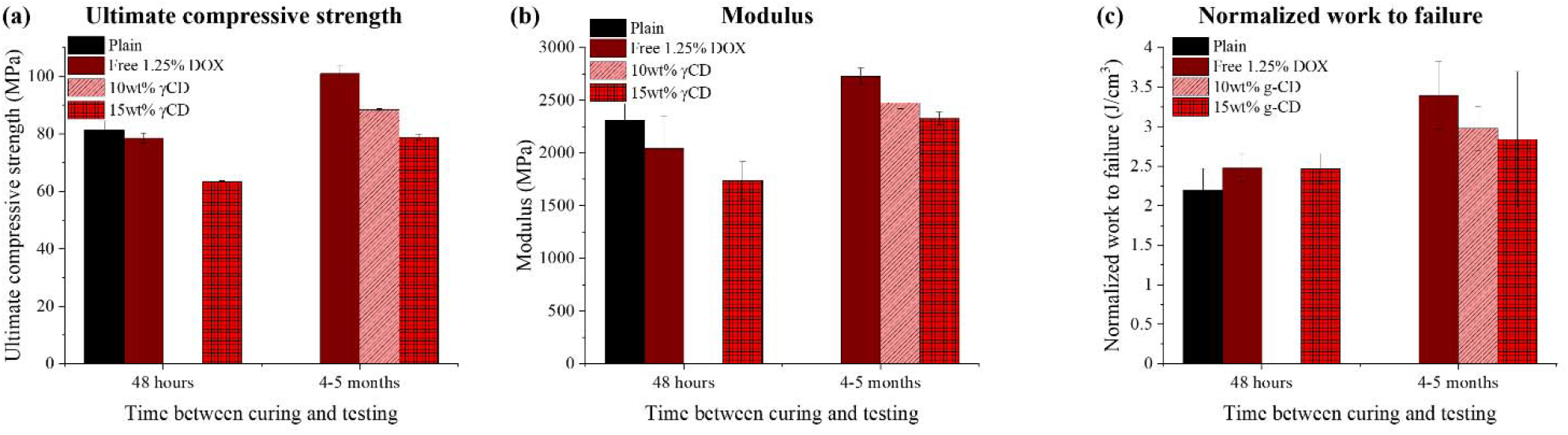
Quantification of ultimate compressive strength (a), modulus (b), and normalized work to failure (c) of PMMA composite cylinders evaluated at either 48 hours (a) or 4-5 months (b) post-fabrication. Sample conditions: plain controls (no DOX, no γ-CD), free 1.25% DOX, and 10wt% and 15wt% DOX-filled γ-CD microparticles in PMMA.

The ultimate compressive strength of the PMMA composites was inversely related to the average pore volume (correlation coefficient = −0.74). Therefore, PMMA composites with a greater pores volume (i.e. PMMA with 15wt% γ-CD) resulted in a weaker mechanical strength. In general, to be used in load-bearing clinical applications (fixation of internal orthopedic prostheses), according to ASTM F451-16 [62] and ISO 5833 standards [65], PMMA bone cement must have a minimum ultimate compressive strength of 70 MPa [66]–[68]. 48 hours post-fabrication, plain PMMA and PMMA containing free 1.25% DOX exceeded the 70 MPa threshold, whereas PMMA containing 15wt% γ-CD fell slightly short of this cutoff (~6.6 MPa). However, as our 4-5 month post-fabrication studies have exhibited, the PMMA continued to strengthen over time as we hypothesize that residual entrapped monomers reacted. Consequently, over time the PMMA composites (i.e. those containing 10wt% or 15wt% γ-CD microparticles) dramatically surpassed the necessary 70 MPa threshold to be used for fixation of internal orthopedic prostheses (10wt% γ-CD = 88.36 ± 0.42 MPa, 15wt% γ-CD = 78.80 ± 1.02 MPa). Furthermore, past studies from our group have demonstrated that preparation technique of the PMMA (i.e. hand- versus vacuum-mixing) can impact the mechanical strength and porosity of PMMA composites [58]. Therefore, techniques such as vacuum-mixing could be investigated to further enhance the compressive strength of the PMMA composites containing γ-CD.

### Quantification of DOX filling of PMMA composites

To evaluate the maximum amount of DOX that could be filled into the PMMA composites, leaching studies were performed. **Table 2** provides a quantification of the average (raw) and normalized mass of DOX initially filled (prefilled) into samples with and without γ-CD microparticles. Free 1.25% DOX (no γ-CD) PMMA samples had over 100-fold more DOX than any of the PMMA γ-CD composites. PMMA containing 10wt% γ-CD had 3.5-fold more DOX than PMMA containing 5wt% γ-CD. Composites containing 15wt% γ-CD had significantly more DOX than PMMA containing either 5wt% (10-fold increase) or 10wt% (2.9-fold increase) γ-CD.

**Table 2.**
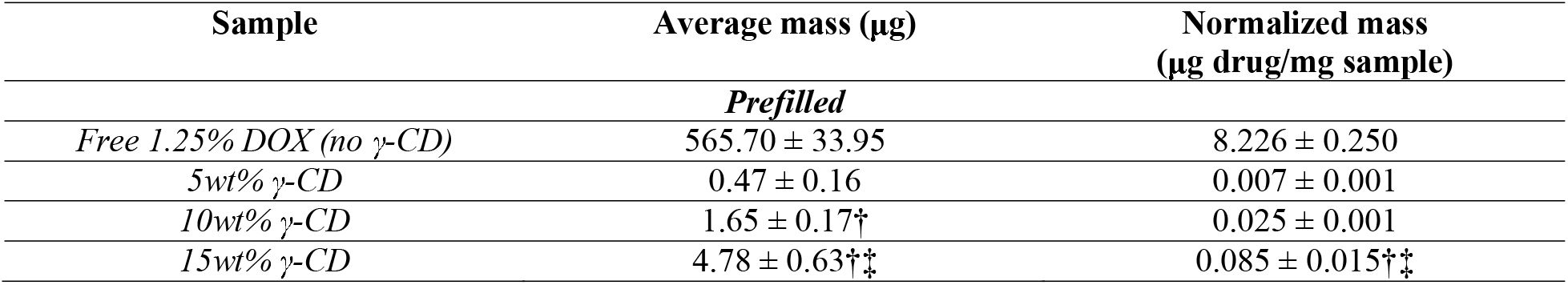
Quantification of initial DOX filling (prefilled) in PMMA γ-CD composites. Statistically significant difference of samples relative to 5wt% γ-CD PMMA (†) and 10wt% γ-CD PMMA (‡).

In conjunction with the studies quantifying the amount of DOX prefilled in PMMA composites, Maestro fluorescence imaging was used to quantify the relative fluorescence intensity of the surface of the DOX-filled PMMA composite samples. **Figure 7a** depicts representative images of PMMA samples without γ-CD (free 1.25% DOX) and containing either 5, 10, or 15wt% γ-CD where the red/orange color indicated the relative intensity of DOX in each sample. Upon visual inspection, there was a gradual increase in the DOX fluorescence intensity as the amount of γ-CD in the PMMA increased. Interestingly, in samples with either 10wt% or 15wt% γ-CD there was an increased DOX fluorescence intensity on the outer periphery. Due to the large content of DOX in the PMMA with free 1.25% DOX, the fluorescence signal was saturated by the system and is not reported.

**Figure 7.**
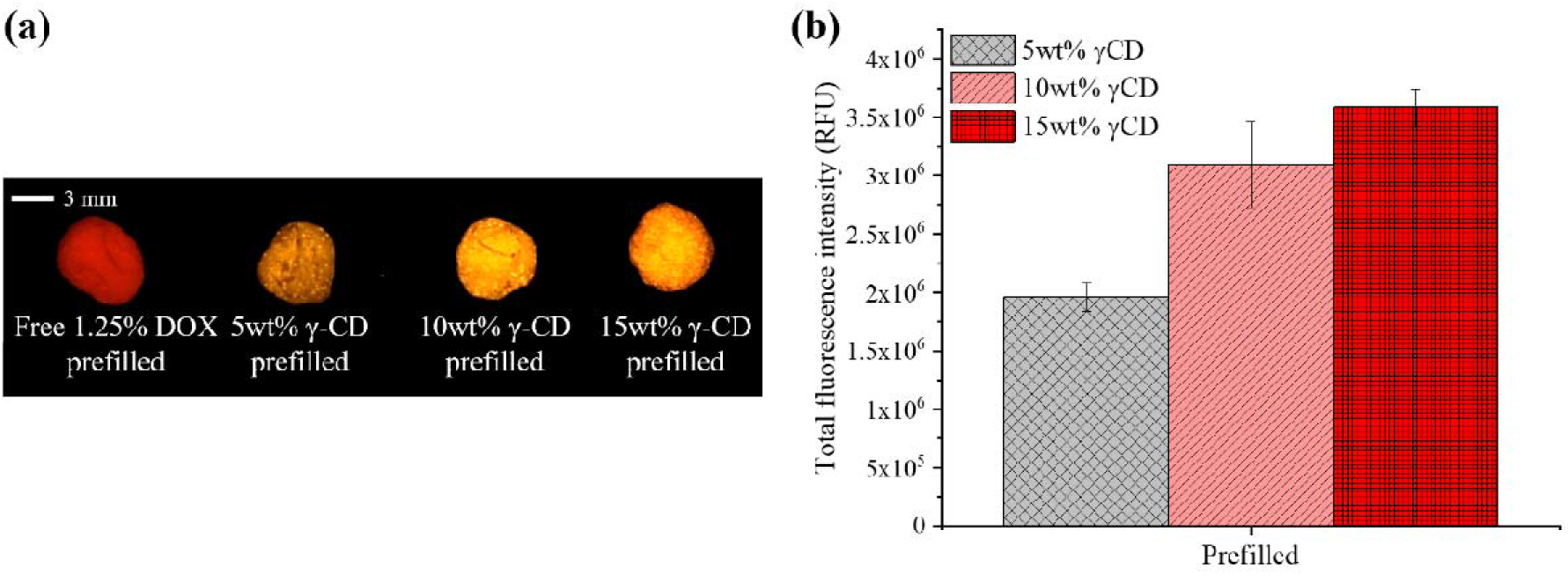
(a) Representative fluorescent images of DOX initially filled (prefilled) into PMMA without γ-CD (free 1.25% DOX) and PMMA with either 5, 10, or 15wt% γ-CD microparticles. Red/orange color was reflective of the distribution and intensity of DOX on the surface of each sample. (b) Quantification of the total fluorescence intensity of the surface of PMMA composites prefilled with DOX.

The intensity of the fluorescence signal of PMMA composite samples prefilled with DOX was quantified and is shown in **Figure 7b**. In accordance with the representative images in **Figure 7a** (and the quantification in **Table 2**), as the quantity of γ-CD in the PMMA increased, there was an increase in the total fluorescence intensity. Samples with 10wt% or 15wt% γ-CD microparticles had a significantly greater DOX fluorescence intensity than those with only 5wt% γ-CD (5wt% γ-CD = 1.96 × 10 ± 1.2 × 10 RFU, 10wt% γ-CD = 3.10 × 10 ± 3.6 × 10 RFU, 15wt% γ-CD = 3.58 × 10^6^ ± 1.6 × 10^5^ RFU). There was no statistically significant difference in the fluorescent signal from samples with either 10wt% or 15wt% γ-CD microparticles.

### Quantification of DOX release from prefilled PMMA composites

After evaluating the total amount of DOX initially filled (prefilled) in the PMMA composites, the DOX release kinetics were evaluated under infinite sink conditions. Due to the strong binding affinity of DOX for γ-CD, a hydrophobic sink (5% Tween 80) was added to the release media (PBS) to assist with the release of DOX. The cumulative and daily amount of DOX released from the PMMA composites is depicted in **Figure 8**. In general, there was significantly more DOX released from PMMA without γ-CD (free 1.25% DOX) than PMMA composites containing 10wt% or 15wt% γ-CD microparticles. Specifically, there was 40.92 ± 3.67 μg DOX cumulatively released from PMMA with free 1.25% DOX, which was nearly 21.4x and 6.8x greater than the amount released from PMMA with 10wt% (1.91 ± 0.18 μg) or 15wt% γ-CD (6.04 ± 0.24 μg), respectively. Interestingly, after 2280 hours (95 days), only ~6% of the theoretical amount of DOX initially filled in PMMA with free 1.25% DOX was released, whereas PMMA containing either 10wt% or 15wt% γ-CD microparticles demonstrated 100% of the theoretical initial amount released after 2040 hours (85 days) and 2424 hours (101 days), respectively. Composites with γ-CD generally had a decreased initial burst release of DOX relative to PMMA with free 1.25% DOX. Specifically, within the first 24 hours, 59.6% of the cumulative DOX released from PMMA with free 1.25% DOX was released. Conversely, only 41.9% and 44.7% of the cumulative DOX released from PMMA with 10wt% or 15wt% γ-CD microparticles was released within the first 24 hours, respectively. As shown in **Figure 8b**, there was generally a greater amount of DOX released from the PMMA containing free 1.25% DOX samples and those containing 10wt% or 15wt% γ-CD at each time point. To elaborate, PMMA samples containing free 1.25% DOX on average released 1-2 μg DOX at each time point, whereas those containing 10wt% or 15wt% γ-CD on average released 0.05-0.5 μg and 0.1-1 μg DOX at each time point, respectively.

**Figure 8.**
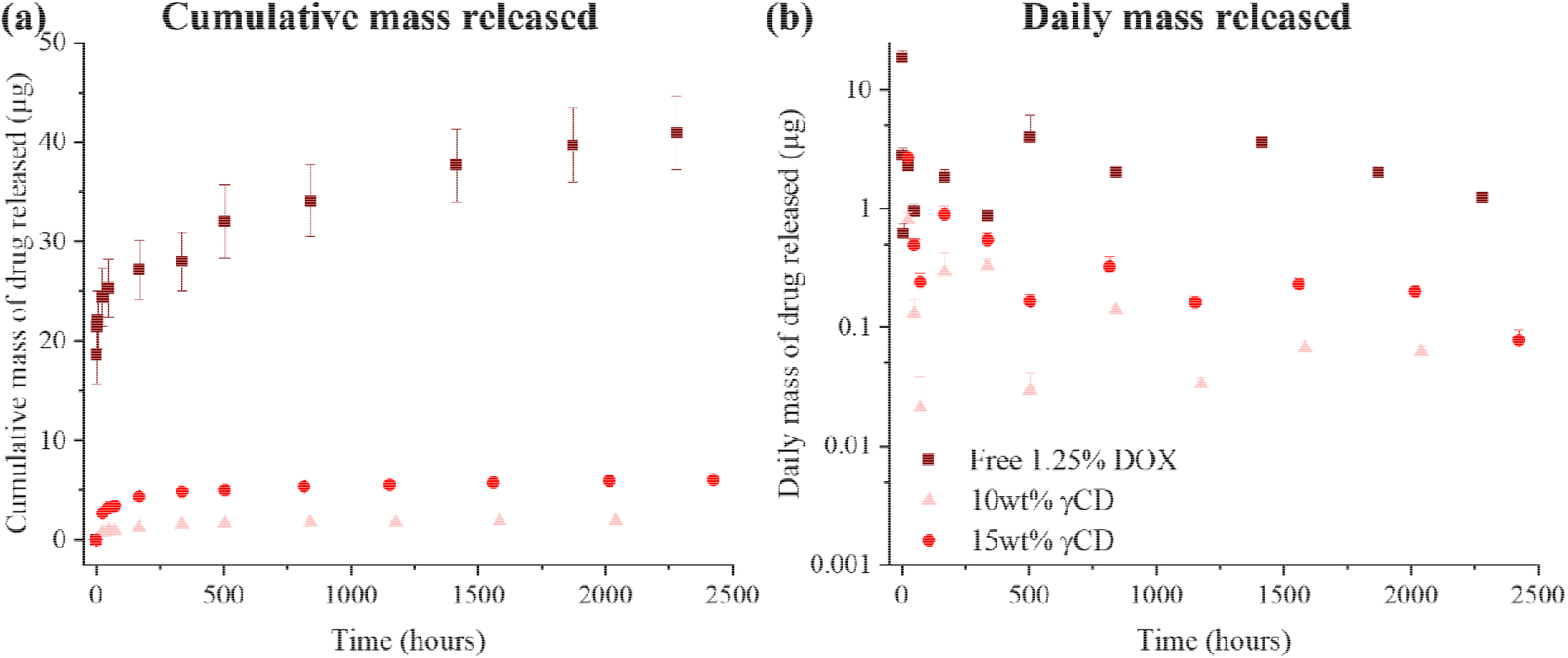
(a) Cumulative and (b) daily mass of DOX released from PMMA composites initially filled (prefilled) with DOX. PMMA had either no γ-CD microparticles (free 1.25% DOX) or 10wt% or 15wt% γ-CD microparticles.

Over a period of approximately 100 days, PMMA containing 15wt% γ-CD prefilled with DOX released > 3-times the amount of DOX relative to PMMA containing 10wt% γ-CD. It was hypothesized that the larger pores in the PMMA containing 15wt% γ-CD enabled the DOX to diffuse out of the PMMA composites more readily. Overall, the cumulative amount of DOX released from the PMMA containing 10wt% and 15wt% γ-CD corresponded with the amount of DOX initially filled (i.e. PMMA containing 15wt% γ-CD had a ~3x greater filling capacity than the PMMA containing 10wt% γ-CD, see **Table 2**). In general, PMMA composites containing either 10wt% or 15wt% γ-CD prefilled with DOX released nearly 100% of the amount initially filled over a prolonged period (~100 days). This was a dramatic improvement over the release kinetics observed with PMMA containing free 1.25% DOX where only ~6% of the initial DOX was released after 95 days. When the majority of the initial DOX remains entrapped in the PMMA, it has the potential to simulate chemoresistance over time if sub-therapeutic doses of DOX are continually released [69]. Therefore, the PMMA composites containing γ-CD may assist in staving off potential chemoresistance with their improved drug release kinetics relative to PMMA containing free 1.25% DOX, however, additional studies are required to investigate this possibility that are outside of the scope of this work. PMMA composites also demonstrated a decreased burst release of DOX initially relative to PMMA containing free 1.25% DOX. Specifically, there was a 25-30% decrease in the percentage of the total DOX released after 24 hours with PMMA composites containing γ-CD microparticles relative to PMMA without γ-CD. Generally, in chemotherapeutic regimens, to reduce off-target toxicities to the patient, it is desirable to have the chemotherapeutic delivered locally in a consistent manner over a prolonged period with a minimal burst release [70], [71].

### Cytotoxicity studies of DOX aliquots from prefilled PMMA composites

To ensure that the amount of DOX released from PMMA composites was therapeutically relevant to kill tumor cells from primary osteosarcoma tumors or potential bone metastases originating from the lungs, a series of MTS cytotoxicity studies were completed against osteosarcoma (MG-63) and Lewis Lung Carcinoma (LLC) cells. In each study, confluent MG-63 or LLC cells were treated with aliquots of DOX in DMEM corresponding to the cumulative amount released from prefilled PMMA composites (free 1.25% DOX, 10wt% γ-CD, or 15wt% γ-CD; see **Table 1**) after 1 day, from 1 to 7 days, and from 7 to 14 days. **Figure 9** depicts the viability of the LLC (a) and MG-63 (b-c) cells incubated with DOX aliquots for either 24 or 48 hours.

**Figure 9.**
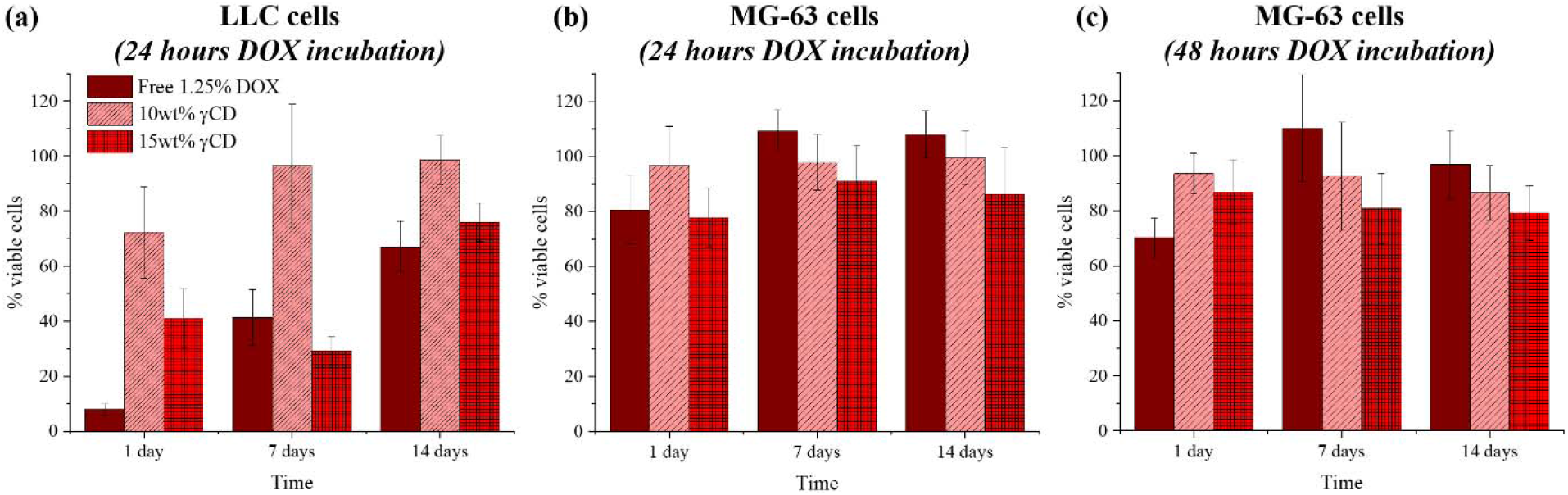
Percent viability of Lewis Lung Carcinoma (LLC) and osteosarcoma (MG-63) cells following 24 hours (a-b) or 48 hours of incubation (c) with DOX aliquots in MTS studies. The concentration of DOX aliquots added to cells was determined based upon the cumulative amount of DOX released from each condition (prefilled PMMA: free 1.25% DOX, 10wt% γ-CD, 15wt% γ-CD) after 1 day, between 1 and 7 days, and between 7 and 14 days.

For the LLC cells, the greatest cytotoxicity was observed using the 1-day cumulative release DOX aliquots from PMMA containing free 1.25% DOX (Free 1.25% DOX = 7.86 ± 1.84% viability, 10wt% γ-CD = 72.23 ± 16.70% viability, 15wt% γ-CD = 41.12 ± 10.65% viability). However, the 7- or 14-day cumulative DOX release aliquots resulted in PMMA containing 15wt% γ-CD demonstrating equivalent or better killing than PMMA containing free 1.25% DOX. Specifically, with the 7-day DOX cumulative release aliquots, the viability of cells with aliquots from PMMA containing free 1.25% DOX was 41.22 ± 10.21%. For DOX aliquots from PMMA containing either 10wt% or 15wt% γ-CD, the viabilities were 96.46 ± 22.34% and 29.21 ± 5.08%, respectively. For the 14-day DOX cumulative release aliquots, the viability of the cells treated with DOX from PMMA with either free 1.25% DOX or 15wt% γ-CD was comparable (Free 1.25% DOX = 67.09 ± 9.26%, 10wt% γ-CD = 98.64 ± 8.85%, 15wt% γ-CD = 75.94 ± 7.12%). It is important to note that with all three cumulative DOX aliquots (i.e. 1-, 7-, and 14-day) the amount of DOX released from PMMA containing 10wt% γ-CD was not sufficient for therapeutically relevant killing as 72.23-98.64% cell viability was exhibited.

For the MG-63 cells, cells were incubated with the cumulative DOX aliquots for either 24 or 48 hours (**Figure 9b-c)**. Generally, there was a slight, but not statistically significant, decrease in the viability of the cells with the longer period of incubation. When the cells were incubated with DOX for 48 hours, DOX aliquots from the PMMA containing free 1.25% DOX exhibited the greatest cytotoxicity with the 1-day cumulative release aliquots. Specifically, when treated with the 1-day cumulative release aliquots from PMMA containing free 1.25% DOX, cells exhibited 70.31 ± 7.01% viability, whereas cells treated with DOX from PMMA containing either 10wt% or 15wt% γ-CD had viabilities of 93.55 ± 7.38% and 86.89 ± 11.43% viabilities, respectively. However, the cumulative DOX aliquots from PMMA containing 15wt% γ-CD significantly outperformed the cytotoxicity of aliquots from PMMA containing free 1.25% DOX at later time points (i.e. 7- and 14-day). With the 7-day cumulative DOX aliquots, cells treated with aliquots from PMMA containing free 1.25% DOX had a viability of 110.13 ± 19.43% and those treated with aliquots from PMMA containing either 10wt% or 15wt% γ-CD had viabilities of 92.67 ± 19.52% and 80.87 ± 12.85%. Similarly, cells treated with 14 day cumulative DOX aliquots from PMMA containing free 1.25% DOX had a viability of 96.81 ± 12.41% and those treated with aliquots from PMMA containing either 10wt% or 15wt% γ-CD had viabilities of 86.61 ± 9.85% and 79.31 ± 9.97%. Similar to the LLC cells, the quantity of DOX released from PMMA with 10wt% γ-CD at each time point (i.e. 1-, 7-, and 14-day) was not therapeutically relevant to kill MG-63 cells with viabilities between 86.61-93.55% following treatment.

Release aliquots obtained from PMMA containing 15wt% γ-CD microparticles appeared to provide the most consistent therapeutic effect over 14 days relative to PMMA containing free 1.25% DOX or 10wt% γ-CD. Cumulative DOX aliquots from PMMA containing 10wt% γ-CD microparticles were not sufficient to provide therapy against either LLC or MG-63 cells. In general, LLC cells demonstrated a greater sensitivity to DOX than MG-63 cells treated with the same DOX concentrations. Since the IC50 value of DOX against MG-63 cells is approximately 0.01 mg/mL, it was not surprising that the DOX aliquots used to treat the MG-63 cells were not therapeutically relevant in most cases [72]. Furthermore, several studies have demonstrated that MG-63 cells can be susceptible to resistance following prolonged exposure to DOX [73], [74]. Therefore, the PMMA composites containing 15wt% γ-CD and DOX may be most amenable for the treatment of bone metastases originating from the lungs, rather than primary osteosarcoma tumors. It is important to note that DOX aliquots from PMMA containing 15wt% γ-CD exhibited greater cytotoxic effects over time than aliquots from PMMA containing free 1.25% DOX. Specifically, against LLC cells, 1-day cumulative aliquots from PMMA containing free 1.25% DOX had a significantly greater cytotoxicity than aliquots from PMMA containing 15wt% γ-CD. However, 7- and 14-day cumulative DOX aliquots from PMMA containing 15wt% γ-CD had comparable or greater cytotoxicity than aliquots from PMMA containing free 1.25% DOX. This was a critical finding because PMMA containing 15wt% γ-CD prefilled with DOX contained significantly less DOX than PMMA containing free 1.25% DOX (~118-times less), yet it still achieved a comparable or improved therapeutic effect for at least 14 days while exposing the patient to a lower dose of DOX without leaving any entrapped.

Monotherapy chemotherapeutics are generally inadequate to eradicate primary osteosarcoma tumors, as was observed in our MTS studies of DOX released from PMMA composites (10wt% and 15wt% γ-CD) against MG-63 cells. Combinatorial chemotherapeutics such as DOX with methotrexate and cisplatin have been shown to more effectively eradicate primary osteosarcoma relative to monotherapy treatments [75]–[78]. Prior work from our group has demonstrated the capacity of PMMA composites to accommodate and release multiple (n = 2) antibiotics in a controlled fashion [56]. Therefore, it is anticipated that the proof-of-concept monotherapy chemotherapeutic PMMA composite system developed in this study could be adapted to accommodate multiple chemotherapeutic agents to improve treatment outcomes of primary osteosarcoma.

### Quantification of refilling and release of DOX from PMMA composites

Upon evaluation of the filling capacity and release kinetics of DOX from prefilled PMMA composites, we were next interested in exploring the possibility of the PMMA composites to be implanted in tissue initially without any drug and later filled (refilled) with DOX via a local bolus injection into the tissue surrounding the implanted PMMA. PMMA composites were embedded in an agarose-based model and refilled with DOX over 48 hours. Results from **Table 3** provide insight into the quantification of the refilling capacity of different PMMA composites with DOX.

**Table 3.**
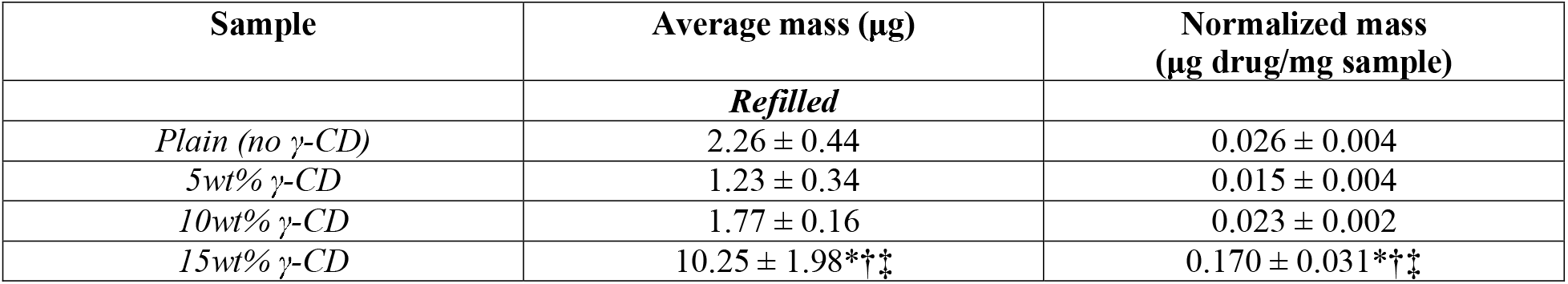
Quantification of DOX refilling into PMMA-γ-CD composites. Statistically significant difference of samples relative to plain PMMA (*), 5wt% γ-CD PMMA (†), and 10wt% γ-CD PMMA (‡).

In the PMMA composites containing γ-CD microparticles, as the content of γ-CD microparticles in the PMMA increased, there was a gradual increase in the amount of DOX refilled.

Specifically, PMMA containing 10wt% γ-CD microparticles refilled 1.5-times more DOX than samples containing 5wt% γ-CD, and PMMA containing 15wt% γ-CD refilled significantly (> 7-times) more DOX than samples containing 10wt% γ-CD. Plain PMMA (without γ-CD) refilled a comparable amount of DOX as PMMA containing 10wt% γ-CD. **Figure 10** depicts representative images of PMMA composites samples before and after refilling with DOX for 48 hours in the agarose model. Upon inspection, in accordance with the values in **Table 3**, plain PMMA refilled with DOX appeared to have a comparable amount of DOX refilled relative to PMMA containing 10wt% γ-CD. Additionally, PMMA containing 15wt% γ-CD was refilled with the greatest amount of DOX as indicated by the red coloration.

**Figure 10.**
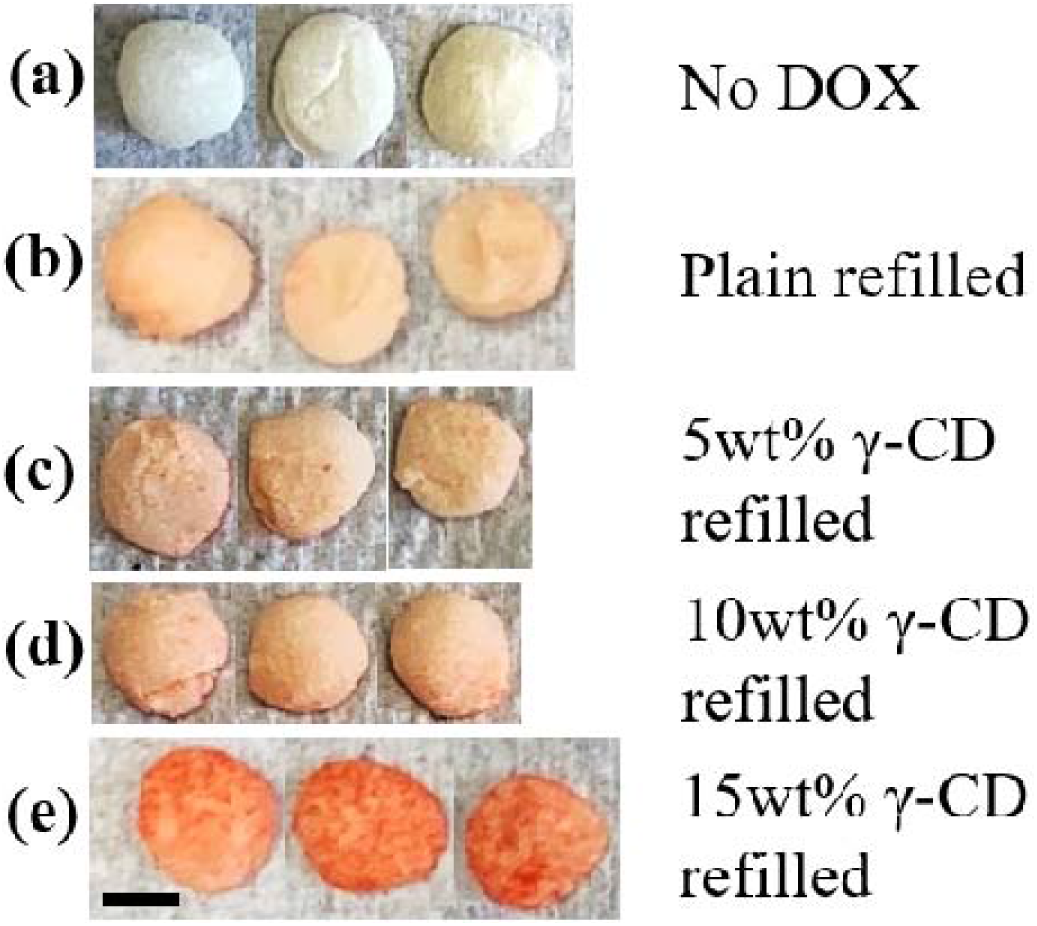
Representative images of PMMA samples before (a) and after DOX refilling for 48 hours in the agarose model (b-e). PMMA samples had either no γ-CD (plain) or 5, 10, or 15wt% γ-CD microparticles. The red/orange color indicated the presence of DOX. Scale bar = 3 mm.

After quantifying the relative amount of DOX refilled into PMMA, we next evaluated the fluorescence intensity of DOX on the surface of the PMMA using Maestro fluorescence imaging.

**Figure 11a** depicts representative images of the fluorescence intensity of DOX refilled into the PMMA composite samples where the top row included controls without any exposure to DOX (empty). As the content of γ-CD microparticles in the PMMA increased, there was a graduate increase in the relative intensity of the DOX fluorescence as indicated by the presence of the red/orange color. Results from the quantification of the total fluorescence intensity of the samples (**Figure 11b**) were in compliance with the qualitative findings. Specifically, there was a statistically significant difference between the fluorescence intensities as the quantity of γ-CD in the PMMA increased (Plain = 1.08 × 10^5^ ± 5.5 × 10^4^ RFU, 5wt% γ-CD = 7.37 × 10^5^ ± 1.9 × 10^4^ RFU, 10wt% γ-CD = 1.14 × 10^6^ ± 6.6 × 10^4^ RFU, 15wt% γ-CD = 2.67 × 10^6^ ± 1.9 × 10^5^ RFU). There was approximately 24.7-times greater intensity of the DOX fluorescence signal on the surface of PMMA containing 15wt% γ-CD relative to plain PMMA. In general, the results from **Figure 11b** followed a pattern like that observed in **Table 3**. Interestingly, results from **Table 3** indicated that there was a comparable amount of DOX refilled into plain PMMA, but the fluorescence studies (**Figure 11**) indicated that plain PMMA had a significantly lower fluorescence than PMMA containing γ-CD. We attribute this discrepancy to the fact that the fluorescence studies only provided information on the amount of DOX filled on the surface of the composites, rather than what had diffused through the entire sample (i.e. **Table 3**).

**Figure 11.**
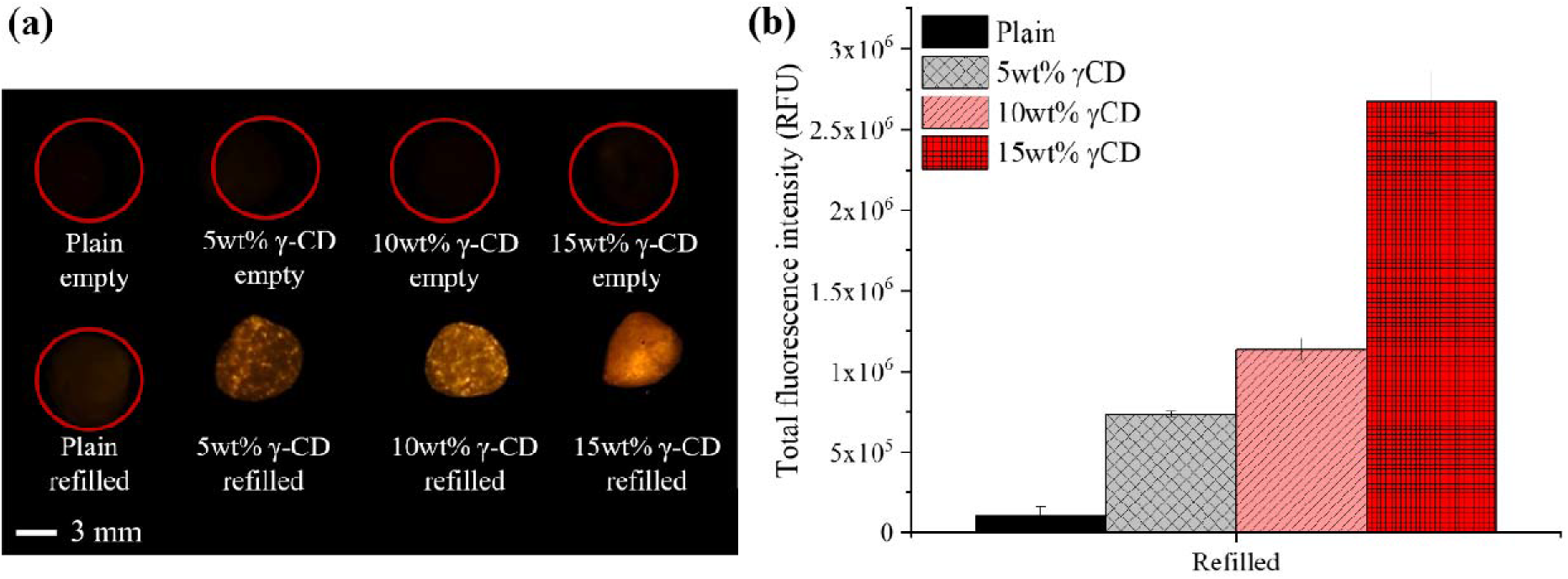
(a) Representative images of the fluorescence intensity of DOX refilled into PMMA composites obtained via Maestro fluorescence imaging. The top row includes control PMMA composite samples not exposed to DOX and the bottom row depicts PMMA samples placed in the agarose refilling model. (b) Quantification of the total fluorescence intensity of PMMA samples refilled with DOX (from bottom row of samples show in Figure 11a).

Following quantification of the DOX refilled into the PMMA composites, the DOX release kinetics were evaluated under infinite sink conditions (PBS with 5% Tween 80) over a period of 74 days. **Figure 12** depicts both the cumulative mass (a) and daily mass (b) of DOX released from refilled PMMA composites with either no γ-CD (plain) or 5, 10, or 15wt% γ-CD microparticles. PMMA with 15wt% γ-CD released about 3-times the cumulative amount of DOX than any of the other compositions (Plain = 4.00 ± 0.37 μg, 5wt% γ-CD = 3.78 ± 0.83 μg, 10wt% γ-CD = 2.68 ± 0.19 μg, 15wt% γ-CD = 11.84 ± 1.09 μg). Additionally, on average there was a greater amount of DOX released at each time point from PMMA containing 15wt% γ-CD than the other compositions (average amount of DOX released at each time point: Plain = 0.2-0.5 μg, 5wt% γ-CD = 0.2-0.5 μg, 10wt% γ-CD = 0.1-0.4 μg, 15wt% γ-CD = 1.0-1.5 μg). Interestingly, within the first 24 hours, only 25.3% of the cumulative drug released from PMMA containing 15wt% γ-CD was released. Alternatively, there was a slightly increased burst release with the other composites with 37.5%, 36.2%, and 55.2% of the cumulative drug released within the first 24 hours from plain and 5wt% and 10wt% γ-CD PMMA, respectively.

**Figure 12.**
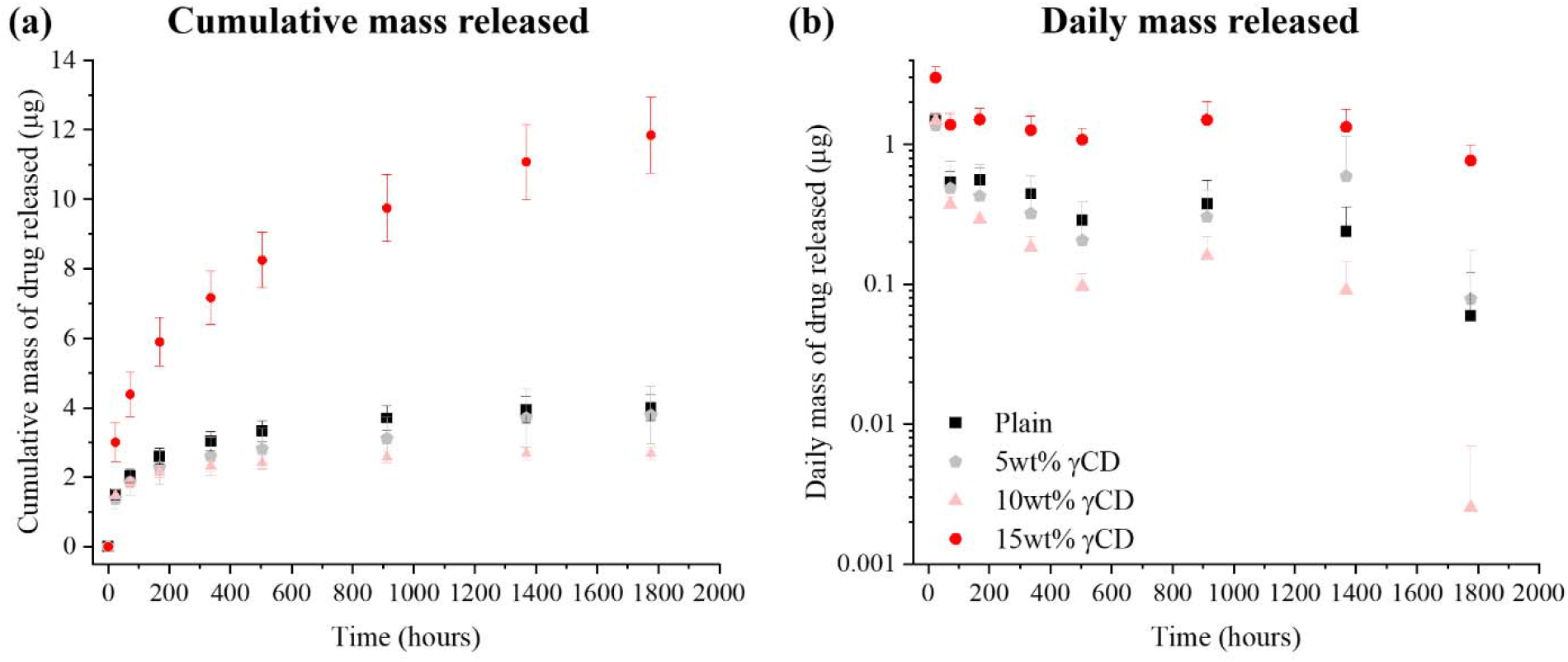
(a) Cumulative and (b) daily mass of DOX released from PMMA composites refilled with DOX in the agarose model. PMMA had either no γ-CD microparticles (plain) or 5, 10, or 15wt% γ-CD microparticles.

In general, PMMA containing 15wt% γ-CD had the greatest DOX refilling capacity. Interestingly, PMMA containing 10wt% γ-CD had a comparable refilling capacity to plain PMMA (without γ-CD) and PMMA containing 5wt% γ-CD refilled less than plain PMMA. These findings were unexpected as the inclusion of any amount of γ-CD microparticles to PMMA was anticipated to increase the DOX refilling capacity relative to PMMA without γ-CD. The discrepancies in the amount of DOX refilled into the composites were likely attributed to the relative amount and distribution of the γ-CD microparticles near the surface of the PMMA that were in direct contact with the DOX during refilling. If the majority of the γ-CD microparticles were clustered in the interior of the PMMA, then in general, less DOX would be refilled into the PMMA. Fluorescent images in **Figure 7a** indicated that a greater amount of DOX appeared to be filled into the periphery of samples containing 15wt% γ-CD, suggesting that perhaps more γ-CD microparticles were at the periphery. Past work from our group has demonstrated that the distribution of γ-CD microparticles in the PMMA can be influenced by the preparation technique of the PMMA (i.e. hand-versus vacuum-mixing) [58]. Therefore, it is possible that the refilling capacity of PMMA composites could be further altered by adjusting the preparation conditions of the bone cement.

A localized and refillable DOX delivery system is highly advantageous for the effective treatment of primary osteosarcoma tumors or bone metastases as it has the possibility of being patient customizable while simultaneously providing long-lasting therapy. Specifically, the maximum amount of drug that can be filled into the PMMA composite can be directly altered through manipulation of the amount and type (i.e. α-, β-, or γ-CD) of CD microparticles incorporated. The rate of drug release from the PMMA composite can be tailored through selection of the type of CD and the relative binding affinity of the drug of interest to the size of the CD pocket. Furthermore, a refillable system offers the potential to implant the PMMA composite initially without any drug and administer a different drug later if the patient requires it through a local bolus injection in nearby tissue. This would help to reduce the patient’s exposure to high concentrations of chemotherapeutics, and in turn, the risk of unwanted side effects. The PMMA composite system is also long-lasting, non-biodegradable, and mechanically strengthens over time and is thereby capable of providing long-term and repetitive therapy.

## Conclusions

In order to develop a proof-of-concept, non-biodegradable, mechanically-robust, refillable depot for the localized and controlled delivery of a monotherapy chemotherapeutic agent (DOX) to either primary or secondary bone lesions, in this study we incorporated insoluble γ-CD microparticles into PMMA bone cement. The PMMA composite system demonstrated the ability to provide a long-term consistent release of DOX with a decreased burst when compared to PMMA without γ-CD. Over a period of ~100 days, 100% of the initial DOX was released from PMMA composites containing γ-CD, whereas only ~6% of the initial DOX was released from PMMA containing free 1.25% DOX over the same duration of time. Furthermore, the PMMA composites demonstrated the capacity to be refilled with additional doses of DOX following implantation, contributing to the patient-customizable nature of the composites. In terms of DOX release kinetics, cytotoxicity, and refilling capacity, PMMA containing 15wt% γ-CD microparticles appeared to be the most suitable composition to yield long-term, consistent, and therapeutically relevant release for treatment of lung-derived bone lesions. While shortly after fabrication (i.e. 48 hours) the compressive strength of PMMA containing 15wt% γ-CD fell slightly short of the 70 MPa threshold for use in fixation of orthopedic prostheses, over time, the composite continued to strengthen and surpassed this threshold. Future work could explore incorporation of combinatorial chemotherapeutics into PMMA composites to improve therapeutic outcomes against primary osteosarcoma tumors.

## Acknowledgements

The authors gratefully acknowledge support through National Science Foundation (NSF) Graduate Research Fellowship Program Grant No. CON50169 (E.L.C.), Case Comprehensive Cancer Center Summer Training for Medical Students (N.K.), Support of Undergraduate Research & Creative Endeavors (SOURCE) (N.Z.), NIH NIAMS Ruth L. Kirschstein NRSA T32 AR007505 Training Program in Musculoskeletal Research (G.D.L.), and NIH R01GM121477 (H.A.vR.). Figure 1 was created using BioRender.com.

## Conflicts of interest

The authors have no conflicts of interest to disclose.

## Notes

### Competing Interest Statement

The authors have declared no competing interest.

